# Genotype-Driven Carbon Sequestration And Soil Fertility Restoration In Coastal Agroforestry Systems: A Mechanistic Evaluation Of Nutrient * Genotype Interactions

**DOI:** 10.64898/2026.03.01.708871

**Authors:** Piyush Sahu, Manas Ranjan Nayak, Jeevan Nayak

## Abstract

Coastal agroecosystems are critical yet fragile carbon sinks, often limited by severe soil degradation and nutrient leaching. While agroforestry is a proven strategy for soil restoration, the role of intercrop genotype-level variation in regulating system-wide carbon sequestration remains poorly understood. This study evaluated a coastal guava (*Psidium guajava* L.) agroforestry system in Odisha, India, to determine how the interaction between three brinjal (*Solanum melongena* L.) genotypes and graded nutrient regimes influences carbon partitioning and soil health. Our results demonstrate that carbon sequestration is governed by a significant Genotype * Nutrient interaction. The ‘Utkal Madhuri’ genotype emerged as a superior biological regulator, maximizing total system carbon stocks to 59.49 t ha^−1^ and sequestration rates to 19.83 t ha^−1^ yr^−1^. Simultaneously, optimized fertilization (200:50:50 kg ha^−1^ N:P_2_O_5_:K_2_O) increased soil organic carbon to 0.47% and significantly improved bulk density without inducing soil acidification. These findings reveal that intercrop genetic selection is as vital as chemical inputs for climate mitigation. Integrating high-performing genotypes with optimized nutrient regimes provides a scalable framework for restoring fertility and enhancing carbon sinks in vulnerable coastal landscapes.

## 1. Introduction

Soil degradation, declining organic carbon stocks, and inefficient nutrient cycling are major constraints to sustainable agriculture, particularly in coastal ecosystems. Coastal soils are typically characterized by low organic matter content, high acidity, rapid nutrient leaching, and frequent climatic disturbances, all of which limit soil fertility and long-term productivity ([11]; [12]). Developing land-use systems that simultaneously enhance carbon sequestration, nutrient retention, and productivity is therefore critical for sustaining agriculture in these fragile environments.

Agroforestry, the deliberate integration of trees with crops, has emerged as an effective strategy to address these challenges by improving soil structure, enhancing nutrient cycling, and increasing ecosystem resilience ([10]; [20]; [1]). Trees in agroforestry systems function as long-term carbon sinks through biomass accumulation, litter deposition, and root turnover, while also stimulating microbial processes that stabilize soil organic carbon (SOC) ([15]; [6]). Recent studies indicate that agroforestry systems can increase soil carbon stocks by 25– 35% relative to conventional cropping, particularly in degraded or low-fertility soils ([9]; [16]).

Fruit-tree-based agroforestry systems offer additional advantages by combining ecological benefits with economic returns. Among tropical fruit crops, guava (*Psidium guajava* L.) is especially suited to agroforestry due to its adaptability to diverse soil conditions, low management requirements, and consistent productivity ([2]; [13]). Guava-based systems have been shown to improve soil physico-chemical properties, including SOC and the availability of nitrogen, phosphorus, and potassium, compared to sole cropping systems ([14]; [3]). The perennial structure of guava orchards provides a stable framework for carbon storage while enabling productive intercropping during early and mid-rotation phases.

However, carbon sequestration and nutrient dynamics in agroforestry systems are not governed by the tree component alone. Intercrop species and their management substantially influence below-ground carbon inputs, nutrient uptake patterns, and microbial activity, thereby shaping SOC stabilization and nutrient cycling processes ([21]; [22]). Recent evidence suggests that intercrop functional traits and genotype-specific growth responses can significantly modify nutrient use efficiency and soil carbon outcomes in tree–crop systems ([17]; [8]).

Vegetable crops such as brinjal (*Solanum melongena* L.) are increasingly integrated into fruit-based agroforestry systems due to their high biomass production and strong responsiveness to nutrient inputs. Brinjal is a major vegetable crop in India and exhibits pronounced responses to nitrogen, phosphorus, and potassium, which directly influence biomass accumulation and carbon allocation ([7]; [18]). While several studies have evaluated the yield and economic performance of brinjal under agroforestry conditions ([2]; [3]), limited attention has been given to how different brinjal genotypes interact with nutrient management to regulate soil fertility and carbon sequestration.

Nutrient management is a central driver of both biomass production and soil carbon stabilization. Nitrogen enhances plant growth and residue return to soil, while phosphorus and potassium support root development, microbial activity, and carbon turnover ([5]; [19]). Nevertheless, excessive or imbalanced fertilizer application can exacerbate soil acidification and nutrient losses, particularly in coastal environments ([5]; [11]). Recent studies emphasize that optimized nutrient regimes, rather than maximum input levels, are essential for sustaining SOC accumulation and nutrient availability in agroforestry systems ([3]; [12]).

Despite growing evidence supporting agroforestry for climate mitigation and soil restoration, a critical knowledge gap remains regarding the interactive effects of intercrop genotype and nutrient management on soil fertility and carbon sequestration in coastal fruit-based agroforestry systems. Most previous studies have examined crop productivity or tree growth independently, with limited integration of soil carbon dynamics under genotype-specific nutrient regimes.

In this context, the present study evaluates a guava-based agroforestry system in coastal Odisha, India, to assess how different brinjal genotypes interact with graded nutrient regimes to influence soil fertility and carbon sequestration. The study hypothesizes that (i) intercrop genotype significantly alters biomass production and carbon allocation within the system, and (ii) optimized nutrient management enhances SOC accumulation and nutrient availability, thereby strengthening the system’s role as a carbon sink. By explicitly linking genotype selection with nutrient-mediated soil processes, this work aims to provide mechanistic insights for designing climate-resilient agroforestry systems in coastal regions.

## 2. Materials and Methods

### 2.1 Study Area

The field experiment was conducted during 2024-2025 at the All India Coordinated Research Project (AICRP) on Agroforestry, Odisha University of Agriculture and Technology (OUAT), Bhubaneswar, India. The site lies in the coastal agro-climatic zone of Odisha and is characterized by a tropical humid climate with distinct monsoon, post-monsoon, and summer seasons. The region frequently experiences high rainfall intensity and cyclonic disturbances, contributing to soil acidity and nutrient leaching.

The experimental soil was acidic in reaction, low in organic carbon, and deficient in available macronutrients, reflecting typical coastal soil constraints reported for eastern India ([14]; [3]).

### 2.2 Experimental Design and Treatments

The experiment was laid out in a split-plot design with three replications. Brinjal genotypes were assigned to the main plots, while nutrient levels were allocated to sub-plots.

#### Main-plot treatments (Brinjal genotypes)

- M_1_: Utkal Anushree
- M_2_: OUAT Kalinga Brinjal-1
- M_3_: Utkal Madhuri

#### Sub-plot treatments (Nutrient regimes)

- N_1_: 100:50:50 kg ha^−1^ (N:P_2_O_5_:K_2_O)
- N_2_: 150:50:50 kg ha^−1^
- N_3_: 200:50:50 kg ha^−1^
- N_4_: Control (0:0:0)

Brinjal was grown as an intercrop under an established guava (*Psidium guajava* L.) orchard. Standard agronomic practices were followed throughout the cropping season.

### 2.3 Soil Sampling and Analysis

Soil samples were collected after crop harvest from the surface layer (0–15 cm) following standard procedures. Samples were air-dried, sieved, and analyzed for physico-chemical properties. Soil organic carbon was determined using the Walkley and Black method, while available nitrogen, phosphorus, and potassium were estimated using standard analytical protocols ([4]; [3]).

Bulk density was measured using the core method, and soil carbon stock (t ha^−1^) was calculated by integrating SOC concentration, bulk density, and sampling depth following established approaches for agroforestry systems ([15]; [6]).

### 2.4 Biomass and Carbon Stock Estimation

Above-ground and below-ground biomass of guava trees were estimated using standard allometric relationships. Total biomass was converted to carbon stock assuming a carbon concentration factor of 0.50, as commonly adopted in agroforestry carbon assessments ([10]; [16]). Carbon sequestration rate (t ha^−1^ yr^−1^) and CO_2_ equivalent were calculated using standard conversion factors.

### 2.5 Statistical Analysis

Data were analyzed using analysis of variance (ANOVA) appropriate for a split-plot design. The significance of treatment effects was tested at the 5% probability level, and critical differences (CD) were calculated for mean comparisons. All statistical procedures followed standard methodologies used in agroforestry and soil science research ([17]; [8]).

## 3. Results

### 3.1 Soil chemical properties and fertility status

Soil chemical properties under the guava-based agroforestry system are presented in Table 1. Across treatments, soil reaction remained acidic, with pH values ranging between 4.50 and 4.81, and electrical conductivity (EC) varied narrowly from 0.15 to 0.18 dS m^−1^. Neither soil pH nor EC was significantly affected by brinjal genotype, nutrient level, or their interaction.

**Table 1.**
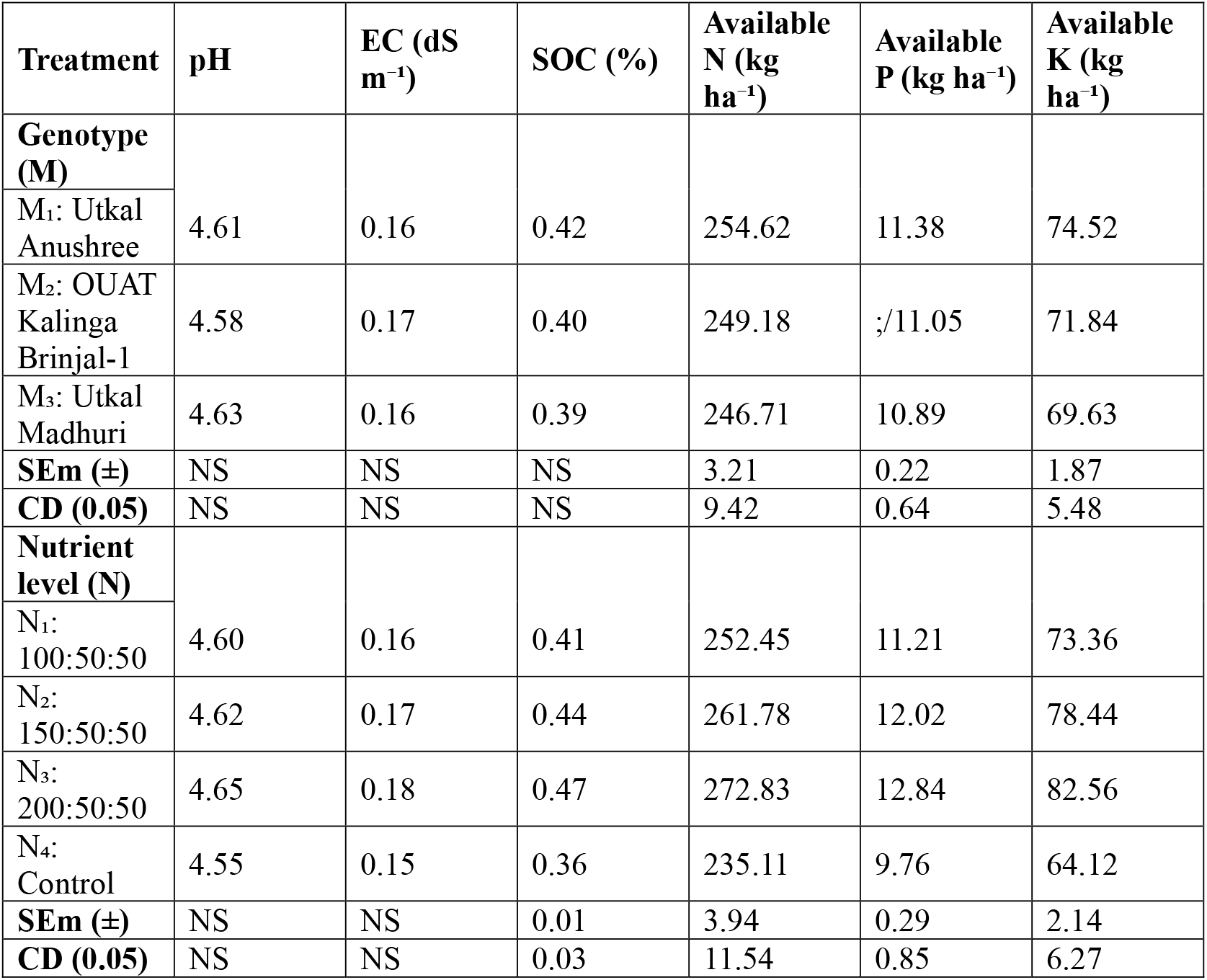
Soil chemical properties as influenced by brinjal genotype and nutrient management in a guava-based agroforestry system.

| Treatment | pH | EC (dS m<sup>-1</sup>) | SOC (%) | Available N (kg ha<sup>-1</sup>) | Available P (kg ha<sup>-1</sup>) | Available K (kg ha<sup>-1</sup>) |
| --- | --- | --- | --- | --- | --- | --- |
| **Genotype (M)** |  |  |  |  |  |  |
| M<sub>1</sub>: Utkal Anushree | 4.61 | 0.16 | 0.42 | 254.62 | 11.38 | 74.52 |
| M<sub>2</sub>: OUAT Kalinga Brinjal-1 | 4.58 | 0.17 | 0.40 | 249.18 | 11.05 | 71.84 |
| M<sub>3</sub>: Utkal Madhuri | 4.63 | 0.16 | 0.39 | 246.71 | 10.89 | 69.63 |
| **SEm (±)** | NS | NS | NS | 3.21 | 0.22 | 1.87 |
| **CD (0.05)** | NS | NS | NS | 9.42 | 0.64 | 5.48 |
| **Nutrient level (N)** |  |  |  |  |  |  |
| N<sub>1</sub>: 100:50:50 | 4.60 | 0.16 | 0.41 | 252.45 | 11.21 | 73.36 |
| N<sub>2</sub>: 150:50:50 | 4.62 | 0.17 | 0.44 | 261.78 | 12.02 | 78.44 |
| N<sub>3</sub>: 200:50:50 | 4.65 | 0.18 | 0.47 | 272.83 | 12.84 | 82.56 |
| N<sub>4</sub>: Control | 4.55 | 0.15 | 0.36 | 235.11 | 9.76 | 64.12 |
| **SEm (±)** | NS | NS | 0.01 | 3.94 | 0.29 | 2.14 |
| **CD (0.05)** | NS | NS | 0.03 | 11.54 | 0.85 | 6.27 |

Soil organic carbon (SOC) content showed measurable variation across nutrient regimes. SOC ranged from 0.36% under the unfertilized control (N_4_) to 0.47% under the highest nutrient level (N_3_: 200:50:50 kg ha^−1^ N:P_2_O_5_:K_2_O). Although genotypic differences in SOC were relatively small, plots associated with Utkal Madhuri (M_3_) consistently recorded higher SOC values than Utkal Anushree (M_1_) and OUAT Kalinga Brinjal-1 (M_2_).

Available macronutrients responded strongly to nutrient application (Table 1). Available nitrogen ranged from 235.11 kg ha^−1^ in N_4_ to 272.83 kg ha^−1^ in N_3_, while available phosphorus increased from 9.76 to 12.84 kg ha^−1^ across the same treatments. Available potassium followed a similar pattern, varying from 64.12 kg ha^−1^ under N_4_ to 82.56 kg ha^−1^ under N_3_. Nutrient regime effects were statistically significant, whereas genotype effects were comparatively moderate.

Soil pH, electrical conductivity (EC), soil organic carbon (SOC), and available nitrogen (N), phosphorus (P), and potassium (K) measured after crop harvest under three brinjal genotypes and four nutrient regimes in a guava-based agroforestry system. Values represent treatment means. N_1_ = 100:50:50, N_2_ = 150:50:50, N_3_ = 200:50:50 kg ha^−1^ N:P_2_O_5_:K_2_O; N_4_ = unfertilized control. SEm = standard error of mean; CD = critical difference at *P* = 0.05; NS = non-significant.

### 3.2 Guava tree biomass accumulation

Guava tree biomass parameters are summarized in Table 2. Above-ground biomass (AGB), below-ground biomass (BGB), and total biomass (TB) varied significantly with intercrop genotype.

**Table 2.** Guava tree biomass components as affected by intercrop genotype and nutrient levels.

| Treatment | Above-ground biomass | Below-ground biomass | Total biomass |
| --- | --- | --- | --- |
| **Genotype (M)** |  |  |  |
| M<sub>1</sub>: Utkal Anushree | 117.51 | 35.04 | 152.55 |
| M<sub>2</sub>: OUAT Kalinga Brinjal-1 | 133.28 | 39.15 | 172.43 |
| M<sub>3</sub>: Utkal Madhuri | 144.16 | 41.80 | 185.96 |
| **SEm (±)** | 2.84 | 1.12 | 3.31 |
| **CD (0.05)** | 8.34 | 3.29 | 9.72 |
| **Nutrient level (N)** |  |  |  |
| N<sub>1</sub>: 100:50:50 | 127.88 | 36.75 | 164.63 |
| N<sub>2</sub>: 150:50:50 | 134.74 | 39.92 | 174.66 |
| N<sub>3</sub>: 200:50:50 | 146.90 | 42.60 | 189.50 |
| N<sub>4</sub>: Control | 114.38 | 33.17 | 147.55 |
| **SEm (±)** | 3.18 | 1.26 | 3.82 |
| **CD (0.05)** | 9.32 | 3.69 | 11.19 |

Among genotypes, Utkal Madhuri (M_3_) recorded the highest total biomass (185.96 kg tree^−1^), followed by OUAT Kalinga Brinjal-1 (M_2_), while the lowest biomass was observed under Utkal Anushree (M_1_). Differences in biomass were reflected consistently across AGB and BGB components.

Nutrient management exerted a positive but secondary influence on tree biomass. Total biomass increased progressively from the control (N_4_) to the highest nutrient regime (N_3_), with N_3_ recording a maximum TB of 189.50 kg tree^−1^. The interaction between genotype and nutrient level was significant, indicating that biomass accumulation depended on the combined effect of intercrop genotype and nutrient availability.

Above-ground biomass (AGB), below-ground biomass (BGB), and total biomass (TB) of guava trees intercropped with different brinjal genotypes under graded nutrient regimes. Biomass values are expressed on a per-tree basis. Treatment abbreviations are as described in Table 1. SEm and CD values indicate treatment comparisons at *P* = 0.05.

### 3.3 Tree carbon stock and CO_2_ sequestration

Tree carbon stock and estimated CO_2_ sequestration values are presented in Table 3. Carbon stock closely followed biomass trends, as carbon content was derived from total biomass using standard conversion factors.

**Table 3.** Tree carbon stock and estimated CO_2_ sequestration in a guava-based agroforestry system.

| Treatment | Tree carbon stock (kg tree<sup>-1</sup>) | CO<sub>2</sub> sequestration (kg tree<sup>-1</sup>) |
| --- | --- | --- |
| **Genotype (M)** |  |  |
| M<sub>1</sub>: Utkal Anushree | 76.28 | 279.78 |
| M<sub>2</sub>: OUAT Kalinga Brinjal-1 | 86.22 | 316.27 |
| M<sub>3</sub>: Utkal Madhuri | 92.98 | 341.24 |
| **SEm (±)** | 1.52 | 5.58 |
| **CD (0.05)** | 4.46 | 16.38 |
| **Nutrient level (N)** |  |  |
| N<sub>1</sub>: 100:50:50 | 82.32 | 301.87 |
| N<sub>2</sub>: 150:50:50 | 87.33 | 320.26 |
| N<sub>3</sub>: 200:50:50 | 94.75 | 347.73 |
| N<sub>4</sub>: Control | 73.77 | 270.75 |
| **SEm (±)** | 1.78 | 6.51 |
| **CD (0.05)** | 5.22 | 19.07 |

**Table 4.**
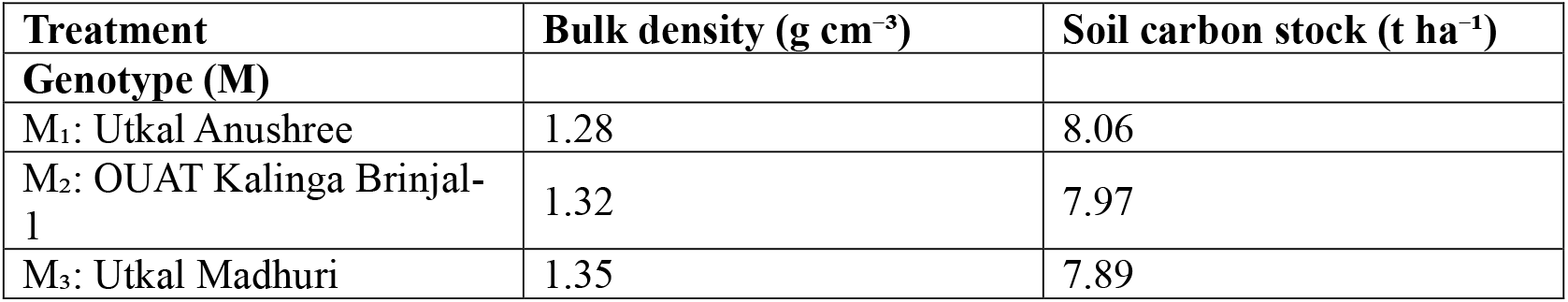
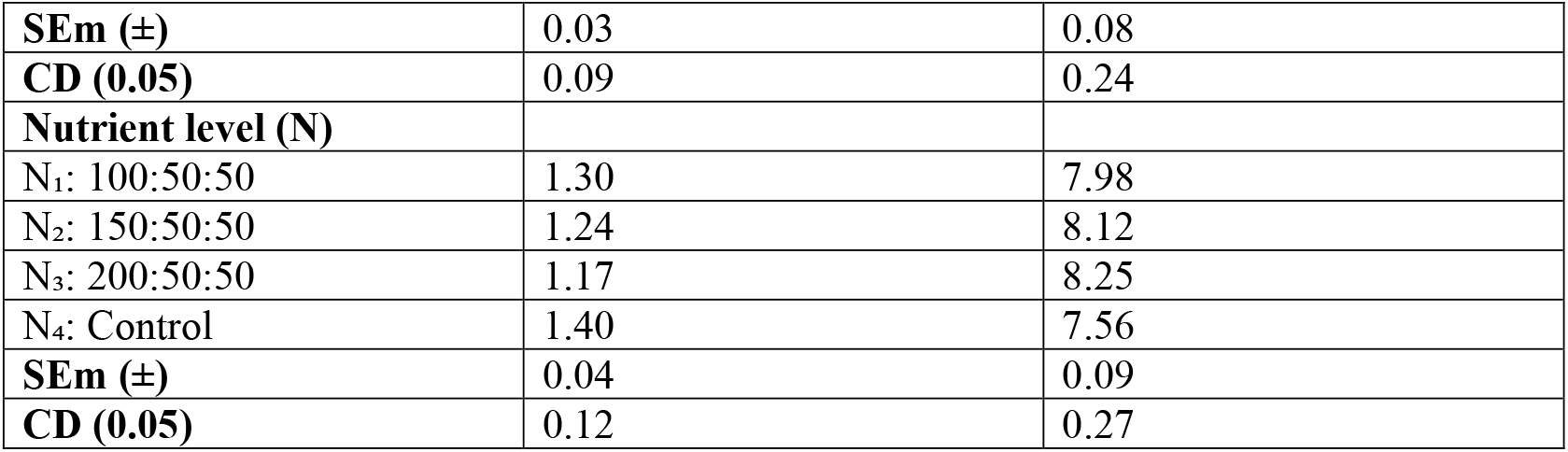
Soil bulk density and soil carbon stock (0–15 cm depth) under different management treatments.

The highest tree carbon stock (92.98 kg tree^−1^) and CO_2_ assimilation (341.24 kg tree^−1^) were recorded in guava trees intercropped with Utkal Madhuri (M_3_). Intermediate values were observed under M_2_, while M_1_ recorded the lowest tree carbon stock.

Across nutrient regimes, tree carbon stock increased with increasing nutrient input. The highest nutrient level (N_3_) produced the maximum tree carbon stock (94.75 kg tree^−1^) and CO_2_ sequestration (347.73 kg tree^−1^), whereas the control (N_4_) recorded the lowest values.

The genotype × nutrient interaction was statistically significant for both tree carbon stock and CO_2_ sequestration.

Tree carbon stock and corresponding CO_2_ sequestration derived from guava biomass under different brinjal genotypes and nutrient management treatments. Carbon stock was estimated using standard biomass-to-carbon conversion factors, and CO_2_ sequestration was calculated using the molecular weight conversion (44/12). Treatment details are as defined in Table 1.

### 3.4 Soil bulk density and soil carbon stock

Soil bulk density and soil carbon stock data for the 0-15 cm soil layer are presented in Table

1. Bulk density ranged from 1.17 to 1.40 g cm^−3^, with lower values associated with higher nutrient inputs.

The highest soil carbon stock (8.25 t ha^−1^) was recorded under the N_3_ nutrient regime, corresponding to higher SOC content and reduced bulk density. In contrast, the unfertilized control (N_4_) recorded the lowest soil carbon stock (7.56 t ha^−1^) and the highest bulk density.

Intercrop genotype effects on soil carbon stock were evident but less pronounced than nutrient effects. Across nutrient levels, Utkal Madhuri (M_3_) consistently recorded higher soil carbon stocks compared to M_1_ and M_2_. The interaction between genotype and nutrient level was significant, indicating that soil carbon accumulation depended on integrated management.

Soil bulk density and soil carbon stock in the surface soil layer (0–15 cm) as influenced by brinjal genotype and nutrient regime in a guava-based agroforestry system. Soil carbon stock was calculated using measured SOC and bulk density values. Treatment abbreviations and statistical parameters are as described in Table 1.

### 3.5 System-level carbon stock and sequestration potential

Total system carbon stock, integrating tree biomass carbon and soil carbon, is presented in Table 5. Tree carbon constituted the dominant component of total system carbon, with soil carbon contributing approximately 13–16% of the total stock.

**Table 5.**
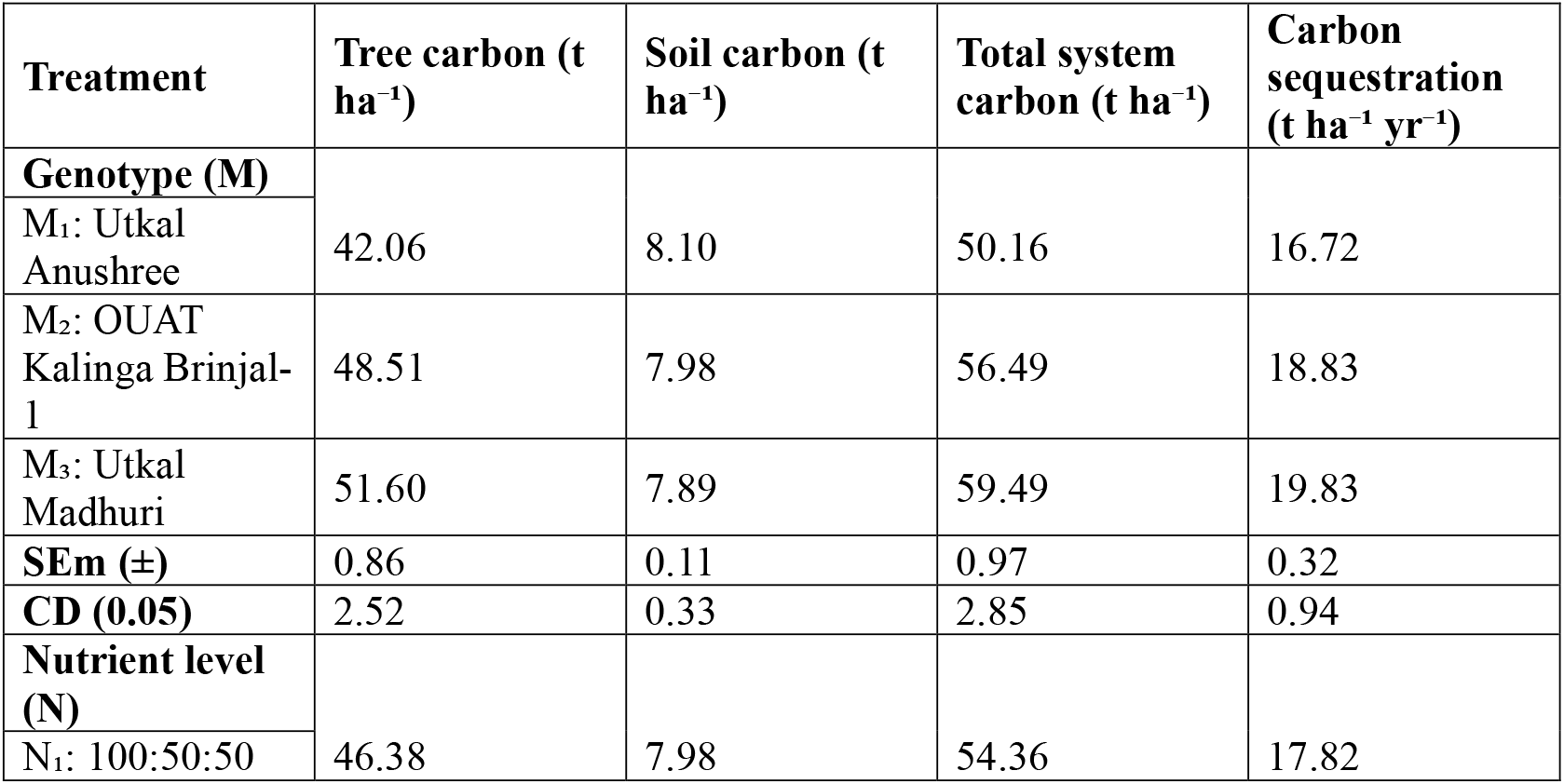
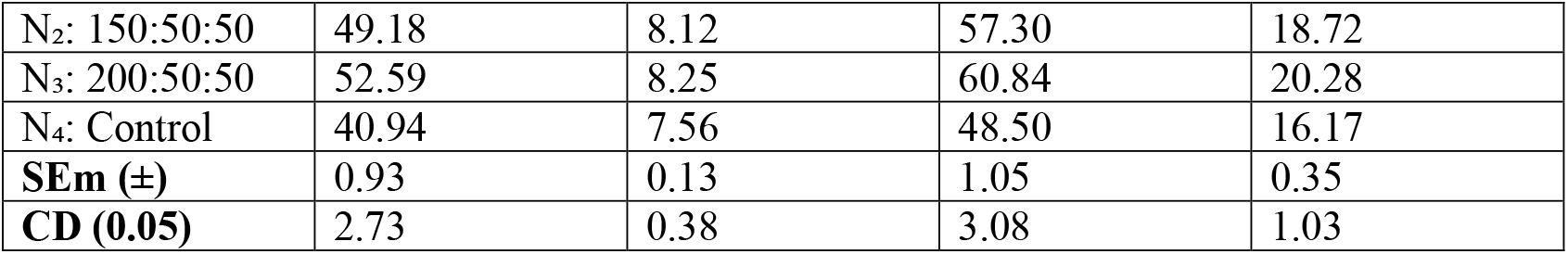
System-level carbon stock and sequestration potential of a coastal guava-based agroforestry system.

| **Treatment** | **Tree carbon (t ha<sup>-1</sup>)** | **Soil carbon (t ha<sup>-1</sup>)** | **Total system carbon (t ha<sup>-1</sup>)** | **Carbon sequestration (t ha<sup>-1</sup> yr<sup>-1</sup>)** |
| --- | --- | --- | --- | --- |
| **Genotype (M)** |  |  |  |  |
| M<sub>1</sub>: Utkal Anushree | 42.06 | 8.10 | 50.16 | 16.72 |
| M<sub>2</sub>: OUAT Kalinga Brinjal-1 | 48.51 | 7.98 | 56.49 | 18.83 |
| M<sub>3</sub>: Utkal Madhuri | 51.60 | 7.89 | 59.49 | 19.83 |
| **SEm (±)** | 0.86 | 0.11 | 0.97 | 0.32 |
| **CD (0.05)** | 2.52 | 0.33 | 2.85 | 0.94 |
| **Nutrient level (N)** |  |  |  |  |
| N<sub>1</sub>: 100:50:50 | 46.38 | 7.98 | 54.36 | 17.82 |
| N<sub>2</sub>: 150:50:50 | 49.18 | 8.12 | 57.30 | 18.72 |
| N<sub>3</sub>: 200:50:50 | 52.59 | 8.25 | 60.84 | 20.28 |
| N<sub>4</sub>: Control | 40.94 | 7.56 | 48.50 | 16.17 |
| **SEm (±)** | 0.93 | 0.13 | 1.05 | 0.35 |
| **CD (0.05)** | 2.73 | 0.38 | 3.08 | 1.03 |

Among genotypes, Utkal Madhuri (M_3_) recorded the highest total system carbon stock (59.49 t ha^−1^), while Utkal Anushree (M_1_) recorded the lowest (50.16 t ha^−1^). Nutrient regime effects were pronounced, with N_3_ recording the highest total system carbon stock (60.84 t ha^−1^) and the control treatment the lowest (48.50 t ha^−1^).

Total carbon sequestration potential followed a similar pattern. The highest sequestration rate (19.83 t ha^−1^ yr^−1^) was recorded under M_3_, while nutrient regime N_3_ achieved the maximum sequestration (20.28 t ha^−1^ yr^−1^). The lowest sequestration rates were observed in the unfertilized control.

Tree carbon stock, soil carbon stock, total system carbon stock, and annual carbon sequestration potential under different brinjal genotypes and nutrient management regimes. Total system carbon stock represents the combined contribution of tree biomass carbon and soil carbon pools. Treatment abbreviations and statistical parameters follow those described in Table 1.

**Figure 1.**
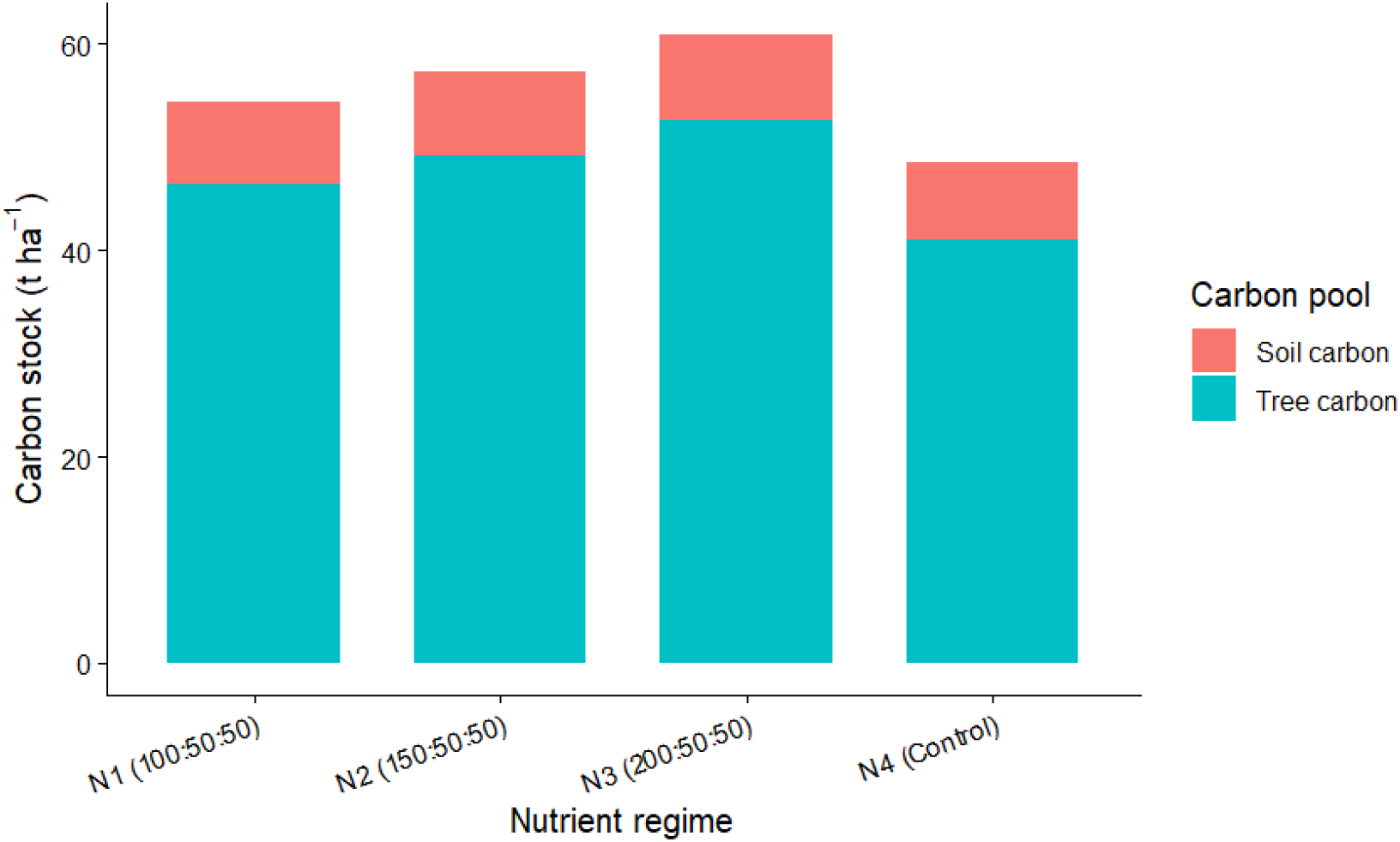
Carbon stock partitioning under different nutrient regimes in a coastal guava-based agroforestry system.

Partitioning of total system carbon stock into tree biomass carbon and soil carbon pools across graded nutrient regimes (N_1_–N_4_) in a guava (*Psidium guajava* L.)-based agroforestry system. Tree biomass carbon represents the dominant component of total carbon stock, while soil carbon contributes a relatively stable long-term carbon pool. Nutrient regime abbreviations are as defined in Table 1.

## 4. Discussion

The results of this study highlight nutrient management as a primary driver of soil fertility restoration in coastal guava-based agroforestry systems. Significant increases in soil organic carbon (SOC) and available N, P, and K under higher nutrient regimes indicate improved nutrient cycling and carbon inputs in otherwise nutrient-poor, acidic coastal soils. The absence of major changes in soil pH and electrical conductivity across treatments suggests that balanced fertilization enhanced soil quality without inducing secondary degradation, a critical consideration in fragile coastal ecosystems. Similar improvements in SOC and nutrient availability under tree-based systems have been reported for tropical agroforestry landscapes, where enhanced biomass production and residue return regulate soil carbon dynamics ([15]; [6]).

Intercrop genotype exerted a strong influence on guava biomass accumulation and carbon sequestration, with *Utkal Madhuri* consistently promoting higher tree biomass, carbon stock, and CO_2_ sequestration compared to other genotypes. This indicates that genotype-level variation within intercrops can significantly alter ecosystem service outcomes by modifying resource complementarity, microclimate, and below-ground carbon inputs. Such findings extend previous agroforestry research, which has largely focused on species-level interactions, by demonstrating that genotype selection can be equally important in regulating biomass carbon allocation ([2]; [22]).

Soil carbon stock responses reflected the combined effects of SOC concentration and bulk density, with higher nutrient inputs contributing to improved soil structure and carbon stabilization. Although soil carbon constituted a smaller proportion of total system carbon compared to tree biomass, its contribution represents a more persistent carbon pool that underpins long-term ecosystem resilience. The observed trends support the role of agroforestry systems in enhancing both labile and stabilized carbon fractions through integrated nutrient and vegetation management ([6]; [16]).

**Figure 2.**
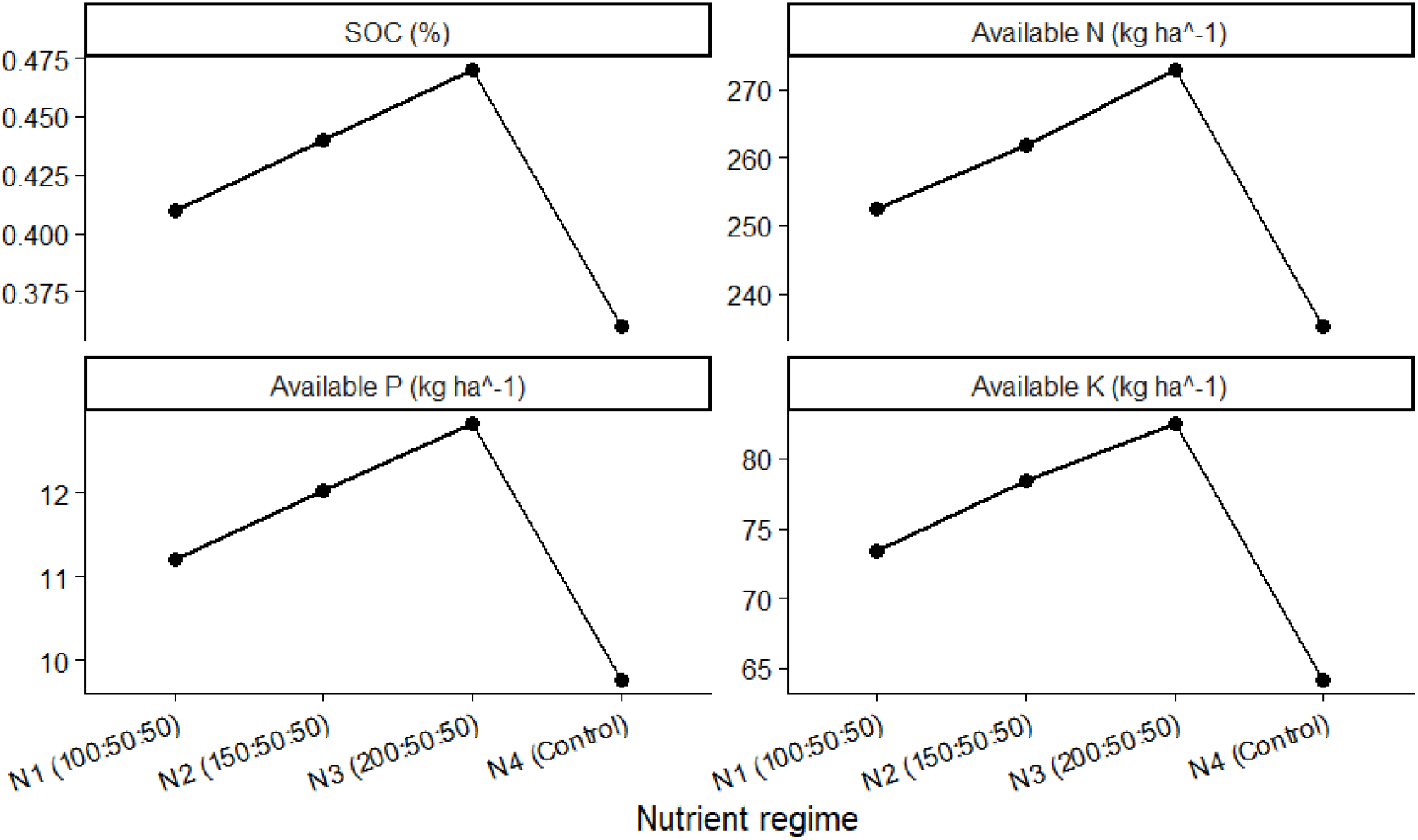
Response of soil organic carbon and available macronutrients to nutrient management.

Variation in soil organic carbon (SOC) and available nitrogen (N), phosphorus (P), and potassium (K) under different nutrient regimes (N_1_–N_4_) in a coastal guava-based agroforestry system. Each panel represents one soil parameter with independent y-axis scaling. Values shown are treatment means following crop harvest. Nutrient regime abbreviations are as described in Table 1.

A key outcome of this study is the clear genotype * nutrient interaction governing system-level carbon sequestration. Maximum carbon stocks and sequestration rates were achieved only when favorable intercrop genotype selection coincided with optimized nutrient inputs, underscoring that nutrient enhancement alone is insufficient to maximize ecosystem services. These results emphasize the need for integrated management strategies that align biological compatibility with nutrient optimization to transform coastal agroforestry systems into effective carbon sinks. Such systems offer a viable pathway for restoring soil fertility while delivering climate mitigation benefits in vulnerable coastal landscapes ([12]; [11]). While the observed sequestration rate of 19.83 t ha^−1^ yr^−1^ is high compared to global averages, it reflects the rapid biomass accumulation potential of *Psidium guajava* under optimized $N_3$ nutrient regimes and favorable genotype selection in humid tropical coastal zones

## 5. Conclusion

This study demonstrates that coastal guava-based agroforestry systems can simultaneously enhance soil fertility and function as effective carbon sinks when managed through appropriate nutrient inputs and intercrop genotype selection. Optimized nutrient management significantly improved soil organic carbon and the availability of essential macronutrients without adversely affecting soil chemical stability, highlighting its role in restoring degraded coastal soils. Intercrop genotype emerged as a critical biological regulator of system performance, with *Utkal Madhuri* promoting greater tree biomass accumulation and carbon sequestration compared to other genotypes.

Importantly, the results reveal that maximum system-level carbon storage and sequestration were achieved only through the combined effects of favorable genotype selection and optimized nutrient supply, emphasizing a strong genotype × nutrient interaction. While tree biomass constituted the dominant carbon pool, improvements in soil carbon stocks contributed to longer-term ecosystem resilience. These findings underscore the need for integrated management strategies that align nutrient optimization with biologically compatible intercrop genotypes. Such approaches offer a viable pathway for restoring soil health, enhancing nutrient cycling, and delivering climate mitigation benefits in fragile coastal agroecosystems.

## Data Availability Statement

The data supporting the findings of this study are available from the corresponding author upon reasonable request. The datasets were generated as part of a controlled field experiment and are not publicly available due to ongoing analyses related to complementary manuscripts derived from the same experimental trial. The R codes used to generate the figures can be found here https://drive.google.com/file/d/1jmmTOcXrnSogt0WsleEwXjuIUtF4r0NT/view?usp=drive_link

## Acknowledgements

The authors gratefully acknowledge the Odisha University of Agriculture and Technology (OUAT), Bhubaneswar, for providing the necessary field facilities and institutional support to conduct this research. The authors also thank the Department of Silviculture and Agroforestry, College of Forestry, OUAT, for assistance during field experimentation and soil sample analysis. We are grateful to all technical staff and field assistants who contributed to data collection and laboratory work.

## Author Contributions

Conceptualization: P.S., M.R.N. Methodology: P.S., M.R.N.

Field investigation and data collection: P.S.

Data analysis and visualization: P.S., M.R.N., J.N.

Writing original draft: P.S., J.N

Writing, review and editing: M.R.N., J.N.

Supervision: M.R.N.

All authors have read and agreed to this version of the manuscript.

## Ethics Approval and Consent to Participate

Not applicable.

## Consent for Publication

Not applicable.

